# Meta-transcriptomic analysis reveals the gene expression and novel conserved sub-genomic RNAs in SARS-CoV-2 and MERS-CoV

**DOI:** 10.1101/2020.04.16.043224

**Authors:** Lin Lyu, Ru Feng, Mingnan Zhang, Qiyu Gong, Yinjing Liao, Yanjiao Zhou, Xiaokui Guo, Bing Su, Yair Dorsett, Lei Chen

## Abstract

**Background:** Fundamental to viral biology is identification and annotation of viral genes and their function. Determining the level of coronavirus gene expression is inherently difficult due to the positive stranded RNA genome and the identification of sub-genomic RNAs (sgRNAs) that are required for expression of most viral genes. In the COVID-19 epidemic so far, few genomic studies have looked at viral sgRNAs and none have systematically examined the sgRNA profiles of large numbers of SARS-CoV2 datasets in conjuction with data for other coronaviruses.

**Results:** We developed a bioinformatic pipeline to analyze the sgRNA profiles of coronaviruses and applied it to 588 individual samples from 20 independent studies, covering more than 10 coronavirus species. Our result showed that SARS-CoV, SARS-CoV-2 and MERS-CoV each had a core sgRNA repertoire generated via a canonical mechanism. Novel sgRNAs that encode peptides with evolutionarily conserved structures were identified in several coronaviruses and were expressed *in vitro* and *in vivo*. Two novel peptides may have direct functional relevance to disease, by alluding interferon responses and disrupting IL17E (IL25) signaling. Relevant to coronavirus infectivity and transmission, we also observed that the level of Spike sgRNAs were significantly higher *in-vivo* than *in-vitro*, while the opposite held true for the Nucleocapside protein.

**Conclusions:** Our results greatly expanded the predicted number of coronaviruses proteins and identified potential viral peptide suggested to be involved in viral virulence. These methods and findings shed new light on coronavirus biology and provides a valuable resource for future genomic studies of coronaviruses.

## BACKGROUND

Corona virus disease 2019 (COVID-19) reached pandemic levels begining March 2020 and brought unprecedented devastation to human lives and the global economy [1]. The causative agent is Severe Acute Respiratory Syndrome – Corona Virus - 2 (SARS-COV-2), a beta coronavirus similar to MERS-CoV, the only other active virulent beta-coronavirus. MERS-CoV is the causative agent of Middle Eastern Respiratory Syndrome (MERS) and is more virulent but less infectious than SARS-CoV-2 and is phylogenetically different from SARS-CoV-2 (less than 90% amino acid sequence homology). Both viruses have a positive single-stranded RNA genome of approximately 30 kilobases that is polyadenylated that encodes 4 structural proteins (spike (S), membrane (M), envelope (E) and nucleocapsid (N)) that play similar roles within each virus. The two viruses diverge with respect to the receptor used for cell entry, their virlulent accessory proteins and the specific function(s) of the 16 non structural proteins (nsp1 to nsp16). Nsp’s are produced by viral proteinase cleavage of two large polyproteins encoded by ORF1a and ORF1b. ORF1 is closest to the 5’ end and is directly translated from genomic RNA upon entrance into host cells and a ribosome skipping mechanism divides it into ORF1a and ORF1b [2]. While MERS-CoV encodes at least 5 accessory proteins (ORF3, ORF4a, ORF4b, ORF5 and ORF8b), SARS-CoV-2 encodes at least 6 (ORF3a, ORF6, ORF7a, ORF7b, ORF8, and ORF10 [3]. All proteins not encoded by ORF1a or ORF1b, must be translated from sub-genomic RNAs (sgRNAs) [4, 5]. SgRNAs are generated via a mechanism termed discontinuous extension that uses short sequences of varying length (usually 6 to 12 nucleotides (nts)) termed Transcription Regulatory Sequences (TRS’s) spaced between genes to pair a 3’ portion of the negative viral strand to a complementary 5’ leader sequence of around 70 nts. This is followed by extension of the negative strand to the 5’ end of the positive strand, generating a short negative strand sgRNA intermediat. The RNA intermediary is then replicated to generate a positive strand sgRNA that encodes viral protein(s) [6] (**Fig. 1A**).

**Fig. 1:**
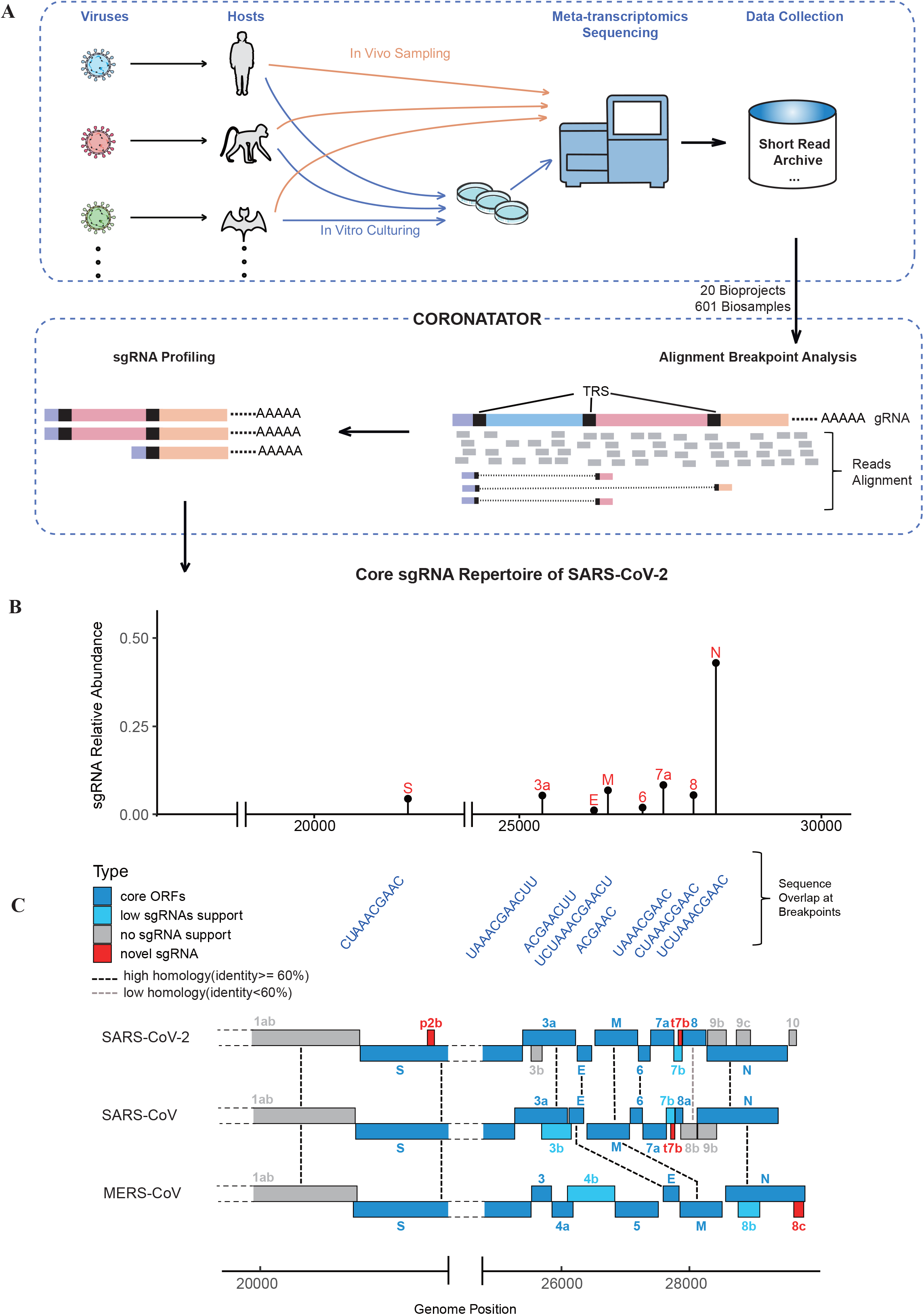
Study overview and the sgRNA profile of SARS-CoV-2. **(A)** Study overview; Top panel, datasets used in this study came from different hosts infected by different coronaviruses, for in-vitro studies, a cell line culturing step was added. These samples were subjected to meta-transcriptomic sequencing and reads were collected from Short Read Archive. Bottom panel, reads mapped to viruses’ genome. Alignment of reads spanning the genome shows breakpoints sites. As viral sgRNAs were formed via the recombination of transcript body and a fixed 5’ leader via TRS homology, breakpoints with 5’ position close to leader TRS were sent to sub-genomic RNA profiling. **(B)** The canonical breakpoints plot of SARS-CoV-2. The ratio of putative sub-genomic RNA, black bar indicates relative abundance of profiled sgRNA. **(C)** ORF annotation and comparison among 3 coronaviruses. Different color of blocks represent sgRNA supportive stat of that gene. Specially, red blocks demonstrate novel sgRNA the algorithm identified.

Annotating viral transcriptomes is fundamental to understanding virus biology, which is a key aspect in combating viral transmission, replication and pathogenesis. Prior coronavirus outbreaks, such as the Severe Acute Respiratory Syndrome (SARS) outbreak in 2003 and the MERS outbreak that began in 2012 and is still ongoing [7, 8], has increased research on these viruses as well as coronaviruses of zoonotic orgin from which human coronaviruses are thought to originate. Comparing transcriptional variation of different coronaviruses may reveal mechanisms behind their distinct pathogenicity and infectivity, and potentially explain the molecular etiology behind how species barriers are crossed. Systematically annotating differences in the transcriptional profiles of virulent coronaviruses that is buried within numerous metatranscriptomic data sets may shed new light on viral transmissibility and virulence. However, even a simple systematic comparison of their *in-vitro* transcriptional profiles is lacking.

For newly emerged SARS-CoV-2 virus, sequencing plays an essential role in diagnosis and monitoring of strain evolution [3, 9]. However, in general, sequencing data sets for SARS-CoV-2 and MERS-CoV are limited to the description of both viral and host transcripts generated during infection of *in-vitro* cell lines as well as model organisms. Analysis of viral transcriptomes orginating from different viral strains in humans is overlooked as suitable analysis tools are lacking.

Sequence homology plays an essential role in the functional annotation of viral genes. However, sequence homology alone does not guarantee protein expression as rapidly mutating RNA viruses can harbor sequence alterations that result in novel or mutated ORFs that are not transcribed nor expressed. Therefore, direct profiling of viral RNAs is the step toward understanding which viral products can actually be generated. For SARS-CoV-2, direct profiling of viral RNAs produced in a cultured cell line was recently conducted using Oxford nanopore technology and identified the existence of a canonical and non-canonical viral transcriptome. All three of these studies used isolated virus strains to infect the Vero cell line isolated from kidney epithelial cells of the African green monkey that does not initiate an interferon (IFN) response upon infection. Although these studies establish a basic characterization of virus transcription, individual studies only characterize viral gene expression of a single viral strain and are unable to determine if viral transcriptional responses are altered in response to even the most basic of immune responses (e.g. IFNγ) [10–12].

We developed a bioinformatics pipeline CORONATATOR (CORONAvirus annoTATOR) to quantify viral gene expression and identify bona-fide sgRNAs in numerous publicly available meta-transcriptomic data sets. Beyond outlining the variation in sgRNA profiles and their relative expression, our analysis identified novel sgRNAs for several different coronaviruses. It also revealed the presence of a core sgRNA repertoire that is shared between SARS and SARS-CoV-2 and one that is unique to MERS-CoV. A subset of novel sgRNAs for SARS-CoV-2 and MERS-CoV appear to be evolutionarily conserved in related coronaviruses found in bat and pangolin. Finally, we show that the transcription of specific sgRNAs differs significantly in-vitro and in-vivo as well as between different coronaviruses.

## RESULTS

### CORONATATOR profiles viral sgRNAs via alignment breakpoint analysis

To systematically identify and compare coronavirus sgRNAs, we sought to identify publicly available coronavirus transcriptomic data sets. As of 2021/09/10, more than 3410427 viral genome sequences were submitted to the Global Initiative on Sharing All Influenza Data (GISAID) [13]. However, few data sets contain the raw sequencing reads. Using the search term “coronavirus” along with manual curation, we located raw reads in a total of 19 Bioprojects within the NCBI Short Read Archive that contain 588 samples for SARS-CoV-2 as well as related coronaviruses, such as SARS and MERS (**Table S1**). We also used an additional dataset with a single sample that was recently published [10].

To profile the sgRNAs present within these data sets, we developed an informatics pipeline (CORONATATOR). CORONATATOR is designed for the utilization of sequences produced by highly accurate second generation sequencing technology that permits identification of TRS sequences from individual reads. Direct RNA Sequencing on the Oxford Nanopore platform can also be used to profile viral sgRNAs but is currently not supported by CORONATATOR due to the limited data set availablility as well as it’s restrictions in terms of sequencing accuracy and read length bias (see **Methods)**.

Briefly, raw reads were first aligned to their respective viral references, i.e. SARS-CoV-2 (GeneBank ID NC_045512.2), SARS-CoV (GeneBank ID NC_004718.3), MERS-CoV (GeneBank ID NC_019843.3) or reference for other species of coronaviruses(**Table S1**). Specific sgRNAs were inferred from alignment breakpoint analysis that identified reads that spanned the junctions between the 5’ leader sequence and more distal genomic sequence (**Fig. 1A and Supplementary Methods**). The relative abundance of a specific sgRNA to all sgRNA’s in a particular sample is analogous to relative gene expression. We constructed a heatmap to determine how viral genotype and viral orgin (e.g.-in-vivo vs in-vitro) influences viral gene expression (**Fig. 3** and discussion below).

CORONATATOR was designed to profile all possible breakpoints. However, to obtain bona-fide sgRNAs, we removed both rare breakpoints and breakpoints that were inconsistent across samples. A complete breakpoint consists of two separate genomic positions (**Fig. 1A**). We also analyzed non-sgRNA breakpoints, for which the 5’ position does not encompass the leader TRS. Our data suggested that non-sgRNA breakpoints are very rare (usually below 0.05% of total sgRNA breakpoints) and inconsistent, as these breakpoints were never identified in more than a single study. We therefore focused on sgRNAs formed with the canonical 5’ leader and a 3’ body part.

### Most predicted coronavirus ORFs can be validated by sgRNA analysis

Many ORFs are annotated for SARS-CoV-2 based on consensus sequence annotation and the existence of some are disputed by proteomics as well as sequencing studies [3, 11]. Only after examination of a large number of data sets from multiple studies were we able to confidently assign commonly annotated ORFs into one of three categories (core, low support and no support) (**Fig. 1B**). Identifying bonafide sgRNAs requires multistudy and multisample analysis as unique artifactual sequences are often generated during sequence library preparation or sequencing([14, 15]). Additionaly, many non-canocical sgRNAs found in low abundance may be random aberrant transcripts without dedicated function (Kim et al, 2020). Therefore, only sgRNAs that are present in multiple studies and data sets are true sgRNA candidates. To classify each viral gene we considered factors such as sgRNA relative abundance, TRS conservation and the potential for leaky ribosome scanning that can be affected by start-codon hijacking[16].

We first validated the commonly annotated ORFs for SARS-CoV-2, SARS-CoV and MERS-CoV by looking at the extent of sequencing evidence that supports the existence of specific sgRNAs. For SARS-CoV-2, thirty four samples were kept after removing those with less than 20 sgRNA reads. To identify robust and consistent sgRNAs that represent the “core” repertoire, which <u>we assign to our first sgRNA catagory</u>, we pooled all sgRNAs identified for a specific virus using a weighted average approach (see Methods) and noted their relative abundance. At a relative abundance of 0.5%, 8 canonical breakpoints emerged corresponding to 8 sgRNA species that harbor 8 well-described ORFs for SARS-CoV-2: S, E, M, N, ORF3a, ORF6, ORF7a and ORF8 (**Fig.1B-C**, **Fig. S2A**, **Table S2, Table S3**). The sgRNA breakpoints for these ORFs are situated between 9 and 162 nt upstream of the start codon. N is the most abundant core sgRNA, representing 54% of the core sgRNAs identified in all samples. The E sgRNA is the least abundant at 1.5%, and the only core protein not identified in recent proteomics studies [11, 17]. ORF7a, M, ORF3a, S, ORF8 and ORF6 are present at 10.6%, 8.4%, 6.9%, 6.1%, 5.9 and 2.7% respectively. Together, these 8 core sgRNAs account for 70% to 100% of the total sgRNAs depending on sample type (e.g. *in-vivo vs in-vitro*), viral strain and read coverage (**Fig. 2**, **Table. S3**).

**Fig. 2:**
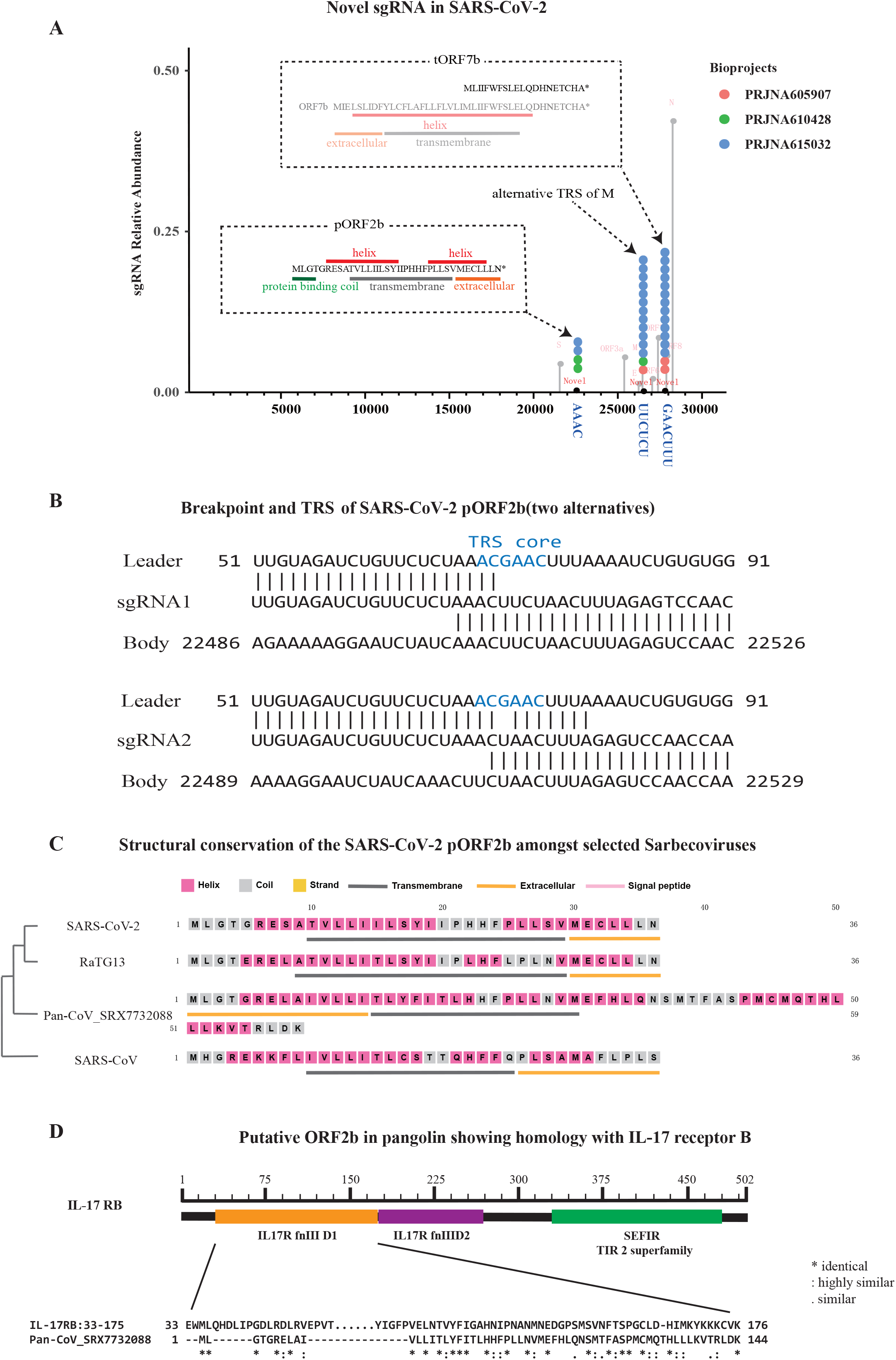
Novel sgRNAs and responsive translational product for SARS-CoV-2. **(A)** Breakpoints plot for SARS-CoV-2 showing the three novel breakpoints at relative abundance cut-off of 0.1%, putative TRS sequences were shown below. Count of classical breakpoints were shown in grey as background. Peptides of novel ORFs, i.e. putative ORF2b (pORF2b) and truncated ORF7b (tORF7b), were shown in inlets, secondary structures of these peptides were predicted and shown in different color. Specially, a complete ORF7b peptide was shown in grey as a reference for the truncated one; **(B)** Sequence homology between leader TRS (top), sgRNA (middle) and body TRS (bottom) for novel sgRNAs, TRS core were shown in blue. **(C)** Structural conservation of novel peptide translated from newly discovered sgRNA putative ORF2b, left panel demonstrates consensus phylogenetic tree of responsive coronavirus determined by genomic sequences, right panel compared second structure of novel peptide predicted by PSIPRED Workbench. **(D)** Putative ORF2b in pangolin CoV shows homology with IL17RB’s fibronectin III like domain, which is also a ligand binding domain.

Beside their high relative abundance, these 8 core sgRNA are also defined by a shared canonical body TRS with a converseved core sequence of “ACGAAC”, which is unique to this group of sgRNAs. This core sequence could be necessary and sufficient for sgRNA formation. Futhermore, the same 8 core sgRNAs, as well the core TRS sequence, were shared by SARS (Fig. S2). The 7 core sgRNAs for MERS following (S, E, M, N, ORF3, ORF4a and ORF5) (Fig. 1C) also utilize this core sequence, with the exception of N that has a TRS which contains “ACGAA”.

A second category of sgRNA was generally present at low relative abundance and does not use this core sequence a conserved core TRS sequence. This category include ORF7b in SARS-CoV-2 and SARS-CoV, ORF3b in SARS and ORF4b and ORF8b in MERS-CoV. For SARS-CoV-2, E has an average relative abundance of 1.5% which is the lowest amoungst the core ones, while ORF7b’s is only 0.02%. This low abundance or low efficiency in sgRNAs formation may result from the use of noncanonical TRSs. This group of sgRNAs do not use the conserved core TRS sequence as core sgRNAs do, meaning the sequence homology they rely on for recombination is always shifted a few bases from the core and quite often they contain mismatches between leader and body TRS.

Other predicted ORFs fell into the third category with no sgRNA support, at least in the data set we examined. When factor in evidence beyond sgRNA support. This category can be futher divided into two sub-categories. The first would be no sgRNA support but can potentially be translated. It has been observed that some coronavirus ORFs can be expressed via a leaky ribosome scanning mechanism [16]. ORF9b of SARS-CoV-2 falls into this sub-category. Indeed, multiple recent proteomics studies showed support for the ORF9b protein product in SARS-CoV-2[17, 18]. Its homolog in SARS-CoV, also named ORF9b, falls in the same category. Interestingly, the ORF7b of SARS-CoV-2 and SARS-CoV were mentioned in previous studies to be in this category (Schaecher et al., 2007), and indeed the long stretch (362 nt in SARS-CoV-2 and 365 nt in SARS) between start codons of ORF7b and preceding ORF7a are void of additional start codons. Yet, these gene products still form their own sgRNAs at low abundance.

The second sub-category would contain the most suspicious ORFs, where sgRNA support cannot be found and intervening start codon between they and the closest sgRNA breakpoint would make their expression very unlikely. This category includes a few commonly annotated ORFs: ORF3b, ORF9c and ORF10 of SARS-CoV-2, ORF8b in SARS-CoV. The several out of frame start codons between these ORFs and preceding ones, along with the absence of corresponding sgRNAs and its absence from proteomic studies [10, 17, 18], strongly argues that these proteins are not generated. Indeed, the existence ORF10 was recently debated in recent manuscripts [11, 12]. The evidence described above indicates the potential pitfalls of conducting experiments on viral products from putative ORFs with no sgRNA or proteomic support. For example, a recent study that generated a synthetic version of the predicted truncated version of ORF3b in SARS-CoV-2 speculated that the putative truncated version in SARS-CoV-2 had a stronger anti-IFN activity than the SARS version [19].

### Identification of novel sgRNAs with non-canonical TRSs in SARS-CoV-2, MERS-CoV and SARS-CoV

As mentioned before, during formation of the core sgRNA repertoire, a body TRS that contains a minimal core sequence will pair with the leader TRS. For each particular core sgRNA, the two TRS’s used must be of the same length and sequence, although the length can vary between sgRNAs (**Fig. 1C**). We found the average length of these canonical TRS’s for SARS-CoV-2 was ~9.6 nts. Interestingly, the same core sequence is used in SARS, while MERS also uses a six nucleotide TRS with a different core sequence (**Fig. S2A,B**).

When we looked for sgRNAs that composed more than 0.2% of sgRNA transcripts, we identified three additional sgRNAs that were present in at least two separate samples and studies (**Fig. 2A**). All three novel sgRNAs contained breakpoints that did not utilize canonical TRS sequences that are present in core sgRNAs. The three breakpoints support the discontinuous extension model of sgRNA formation, as the sequence from the body strand was found in the TRS sequences of the final transcript (**Fig. 2B**, **Fig. S3A-C**). On a separate note, sequence analysis of stranded RNA library preps identified the presence of negative strand sgRNAs, which were not described in the previous Nanopore sequencing manuscripts [10–12]. As previously noted for artificial TRS’s, analysis of these non-canonical breakpoint sequences revealed that TRS’s without perfect complementarity may pair, and/or that large regions of complementarity around a core TRS between the body to itself,maybe used for the formation of sgRNAs (**Fig. 2B**,). Our analysis confirmed that TRS sequences can vary significantly between distantly related viruses and find that canonical TRS sequences can be more than 30 nt in length in some coronaviruses (**Fig. S2D**).

The three novel TRS’s generated three novel sgRNAs that we have termed putative ORF2b (pORF2b), alternative M (aM) and truncated ORF7b (tORF7b). The longest novel sgRNA, pORF2b, is within the S gene and has two alternative TRS’s positioned around 22501. Interestingly, it encodes a novel peptide that has a domain structure that is conserved in closely related coronaviruses, with at least one virus harboring and extended ORF (**Fig. 2B,C**). The second novel breakpoint is located at 26494, 31 nt downsteam of the canonical breakpoint for M. The sgRNA would support M expression, but with an alternative 5’ UTR (**Fig. S3A**). The shortest of the three novel sgRNA’s has its breakpoint positioned at 27761 and codes for a truncated version of ORF7b (tORF7b). The truncation removes the extracellular domain and 14 of the 24 amino acids that comprise the transmembrane domain (**Fig. 2A**, **Fig. S3B**). This sgRNA is expressed at relatively high levels both *in-vivo* and *in-vitro* and likely harbors novel functions (see discussion below).

Translation of pORF2b results in a 36 amino acid peptide. It was predicated by PSIPRED [20] to have a intracellular protein binding coil and two short alpha-helixes that overlap a transmembrane domain, with the second alpha helix partially extracellular (**Fig. 2A, C**). pORF2b was present in 4 samples in two separate studies. The highest expression of pORF2b was observed in a patient derived sample from Washington State in the US (SRX7884411), where it accounted for a substantial 11.1% of the total sgRNAs. In a separate patient sample (SRX7884409) from the same bioproject, the novel ORF represented 1.1% of the sgRNAs identified (**Table. S3**). The virus strains infecting these two patients differed by one nucleotide. Five other patient samples from the same study with different viral strains (**Table. S3**) did not yield sgRNAs for pORF2b. The low breakpoint read numbers for these samples as well as viral strain may contribute to the variable detection of pORF2b *in-vivo*. This indicates that the level of pORF2b transcripts maybe loosely correlated with viral strain and further demonstrates that samples within this bioproject are not cross contaminated with an artifactual pORF2b sgRNA. SgRNA pORF2b was also identified in a separate study (PRJNA615032), in two in-vitro samples that used a different viral strain than any of those identified in the *in-vivo* study (**Table. S3**).

We searched for sequence conservation of pORF2b in other related Sarbecoviruses, including SARS-CoV, HKU3 (bat coronavirus), RaTG13 (a bat coronavirus proposed to be directly related to SARS-CoV-2) and a coronavirus infecting pangolin (SRX7732088)[21]. A corresponding ORF was identified in all four viruses, with the highest level of homology found in RaTG13, with 91.89% nucleotide identity (**Table S4 & Fig. 2C**). Interestingly, pORF2b and more so the pangolin version which has a C terminal extension, share high similarity with the ligand binding domain of human IL17RB **(Fig. 2D and see section “Disscussion”**).

The third novel breakpoint was located at position 27761, within ORF7b, and encodes a truncated version of ORF7b (tORF7b). We identified this transcript and its relative abundance *in-vivo* and *in-vitro* in two separate bioprojects that included more than one viral strain. This transcript was also recently identified in a VERO cell line infected by a single viral strain [10]. Interestingly, this novel sgRNA was expressed at relatively high levels both *in-vivo* and *in-vitro* (**Fig. 3**), and a SARS-CoV homolog of this sgRNA was also present in several samples across two studies. This truncated version of ORF7b is missing the intracellular domain and more than half of its transmembrane domain, while retaining its hydrophilic extracellular domain (**Fig. S3D**). ORF7b is present in the SARS-CoV virion particle and is homologous ORF7b encoded by SARS-CoV-2 [16]. The portion of ORF7b encoded by tORF7b is highly conserved in SARS (**Table S4 & Fig. S3D**). Intrigingly, a previous study observed that a 45 nt deletion in SARS ORF7b that removes much of the transmembrane domain lost in tORF7b, attentuated the induction of interferon-beta, provides a replicative advantage *in-vitro* and *in-vivo* as well as to cells pretreated with interferon-beta [22]. Future research will reveal if this novel sgRNA encodes a novel virulent peptide that has function(s) antagonistic to IFN while subverting the initation of an interferon response.

**Fig. 3:**
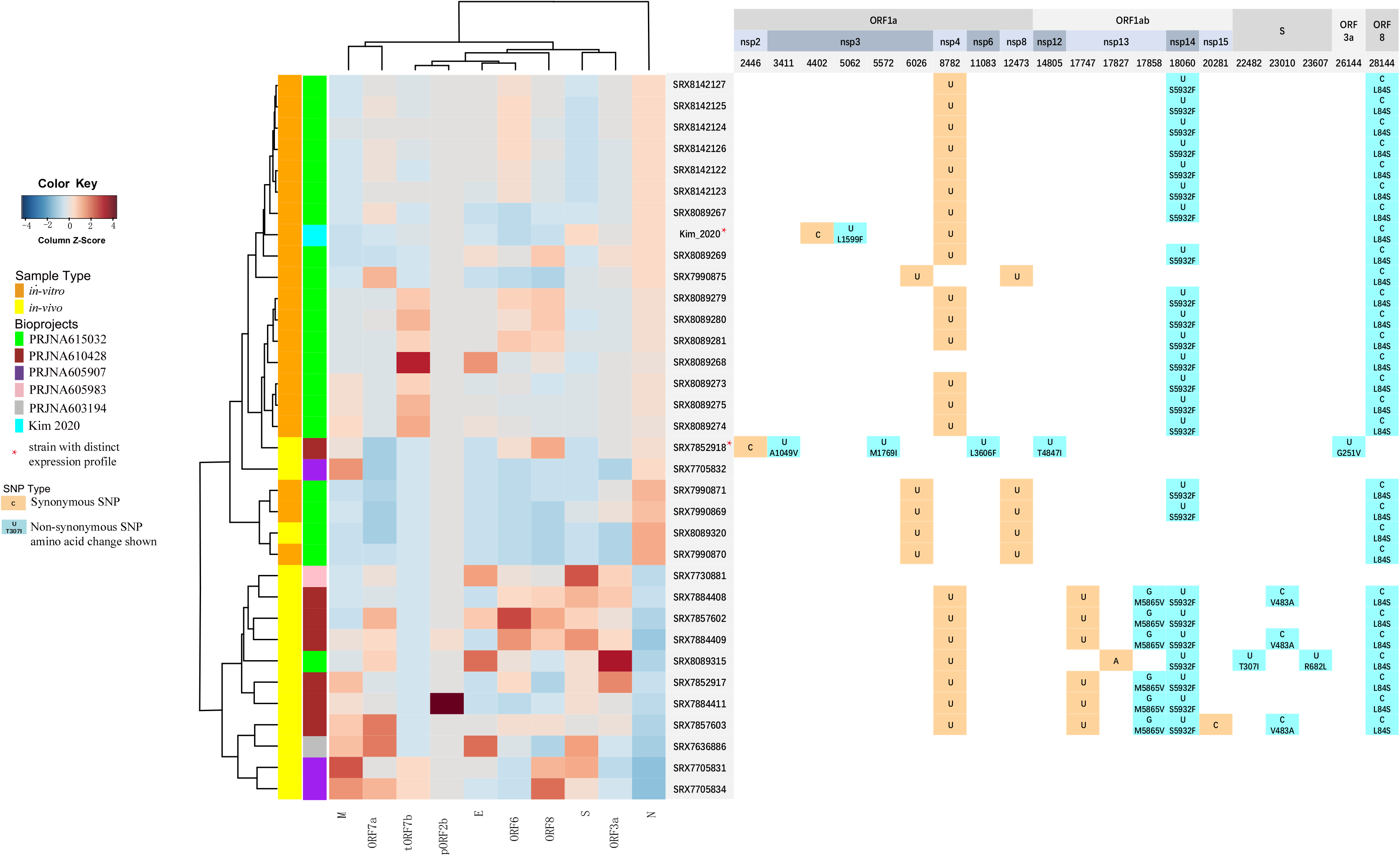
Heatmap of sgRNA expression profile of SARS-CoV-2 with SNP annotation. Left panel shows sgRNA expression profile of SARS-CoV-2 in the transcriptomic or meta-transcriptomic dataset profiled. Right panel shows all the SNP sites with annotation of the responsive biosamples. Interestingly, virus strains from SRX7852918 and Kim et al had distinctive SNP pattern as well as characteristic expression profiles.

We also obtained a significant amount of *in-vivo* and *in-vitro* sequence data sets for MERS-CoV, allowing us to identify abundant non-canonical sgRNAs (**Fig. S2B**). This novel sgRNA (putative ORF8c or pORF8c), is predicted to encode a ORF that translate into a novel 51 amino acid peptide. This novel sgRNA was identified in 5 separate studies, both *in-vivo* and *in-vitro*, ranging in abundance from 0.03% to 1.0% of total sgRNAs. PSIPRED suggest this novel peptide has a transmembrane domain connected to a cytoplasmic helix domain. We also looked for its conservation in other Merbecoviruses, including HKU4, HKU5 and an Erinaceus coronavirus. pORF8c could be found in all 3 with varying conservation (**Fig. S3E**, **Table. S4**). The cytoplasmic N terminal was the most conserved across Merbecovriuses and C terminal elongated versions were observed in HKU5 and Erinaceus (**Fig. S3E**).

To exhaust our search for novel sgRNAs, we lowered our threshold value to a relative abundance of 0.01%, while maintaining our other criteria. This analysis identified additional novel sgRNAs that appeared in more than one study for SARS-CoV-2, SARS-CoV and MERS-CoV (**Table S5**). Additional sequencing and future experiments will determine the significance of pORF2b, tORF7B and aM as well as the numerous other novel sgRNAs present at extremely low abundance.

### CORONATATOR detects experimentally induced alteration of novel pORF8c relative abundance

We next wished to validate the experimental utility of our pipline and validate that a novel sgRNA responds to experimental stimuli in a manner similar to other established viral genes. To accomplish this, we utilized an experimental data set that tested the effects of Gleevec and IFN-β on host gene expression during treatment of MERS-CoV infection in-vitro (PRJNA233943 & PRJNA233944) (**Fig. S5**). Specifically, we analyzed the effects on viral load, viral gene expression and the expression of novel pORF8c. Initial analysis demonstrated that decreased viral load broadened the expression of individual viral genes (**Fig. S5**). Importantly, even at low viral loads, the ratio of N to S remained high, demonstrating that this ratio is not influenced by viral abundance, but by *in-vitro* and *in-vivo* context. The effect of both Gleevec and IFN-β on viral gene expression was not uniform, having different effects on different viral genes. Interestingly, the expression profile of novel pORF8c followed the same trend of N and E with respect to viral load in response to IFN-β and Gleevec. This demonstrates that pORF8c has the same biological response in terms of gene expression as some “core” sgRNAs in this context.

### The relative abundance of Spike sgRNAs is elevated for SARS-CoV-2 *in-vivo*

When processing the data sets, we noticed two distinct patterns of read coverage along the SARS-CoV-2 reference genome that suggested that viral reads originate from two sources. Upon further examination, it was revealed the two sources were in-vivo and in-vitro derived samples (**Fig. S1**). The former is composed of extracellular virion particles and infected host cells present in BALF (human) and nasal washes (Ferret) or lung homogenate (MERS), while the latter is composed of infected cells that are not subject to systemic or sometimes innate (e.g. VERO cells do not produce IFN) anti-viral reponses. *In-vivo* derived viral sequences obtained primarily from BALF for SARS-CoV-2 (primarily BALF) generally covered the entire viral reference length, with little bias towards the sgRNA containing 3’ end. In contrast, highly elevated coverage at the 3’ end of the viral genome was observed in the in-vitro samples due to the formation of nested sgRNAs during viral transcription.

SARS-CoV-2 and MERS-CoV are the only active virulent coronaviruses and are present in both *in-vivo* and *in-vitro* derived metatranscriptomic data sets. We analyzed the relative abundance of sgRNAs generated *in-vivo* and *in-vitro* for both SARS-CoV-2 and MERS-CoV. When comparing the relative abundance of viral sgRNAs generated *in-vivo* to those generated *in-vitro*, it was evident that the ratio of S sgRNAs to N sgRNAs was significantly higher *in-vivo*, especially for SARS-CoV-2 (0.04 in-vitro vs 0.69 in-vivo for SARS-CoV-2, p value 0.0012 with Wilcoxon ranksum test) (**Fig. 4A-B**, **Fig. S4**). The differences in environemental pressures that influence the requirement for these sgRNAs for viral replication, provdes a general explanation for this striking variation in sgRNA levels. The selective pressures may alter viral transcriptional responses that promote viral propogation. For example, the primary function of the S protein centers around host cell recognition and invasion while the primary function of the N protein centers around the regulation of viral RNAs to promote viral replication. This is mediated by direct binding of the 3’ end of the viral genome, the viral packaging signal as well as TRS’s [23–25]. Another explanation, which is not mutually exclusive, is that the increased S/N ratio *in-vivo* is due to an altered viral Replication Transcription Complex (RTC) that favors TRS read-through, preferentially generating longer sgRNAs. Such a “global” alteration of viral transcription likely involves host factors, as observed for Infectious bronchitis virus (IBV), a gamma coronavirus in which the N protein in phosphorylated by cellular GSK3 to recruit the helicase DDX1 to promote TRS read through during the formation of long sgRNAs [26]. In this regard, the N protein is generated from the shortest sgRNA while the S protein is generated from the longest. Future electron microscopy studies on *in vivo* and *in-vitro* viron particles will determine if Spike sgRNA abundance in SARS-CoV-2 correlates with spike protein levels on viron surfaces. Other examples of sgRNAs that are significantly differentially expressed in-vitro and in-vivo include the overall increase in the levels of accessory sgRNAs that act via multiple pathways to quell the immune response in both SARS-CoV-2 and MERS-CoV (**Fig. S4**, [27]).

**Fig.4:**
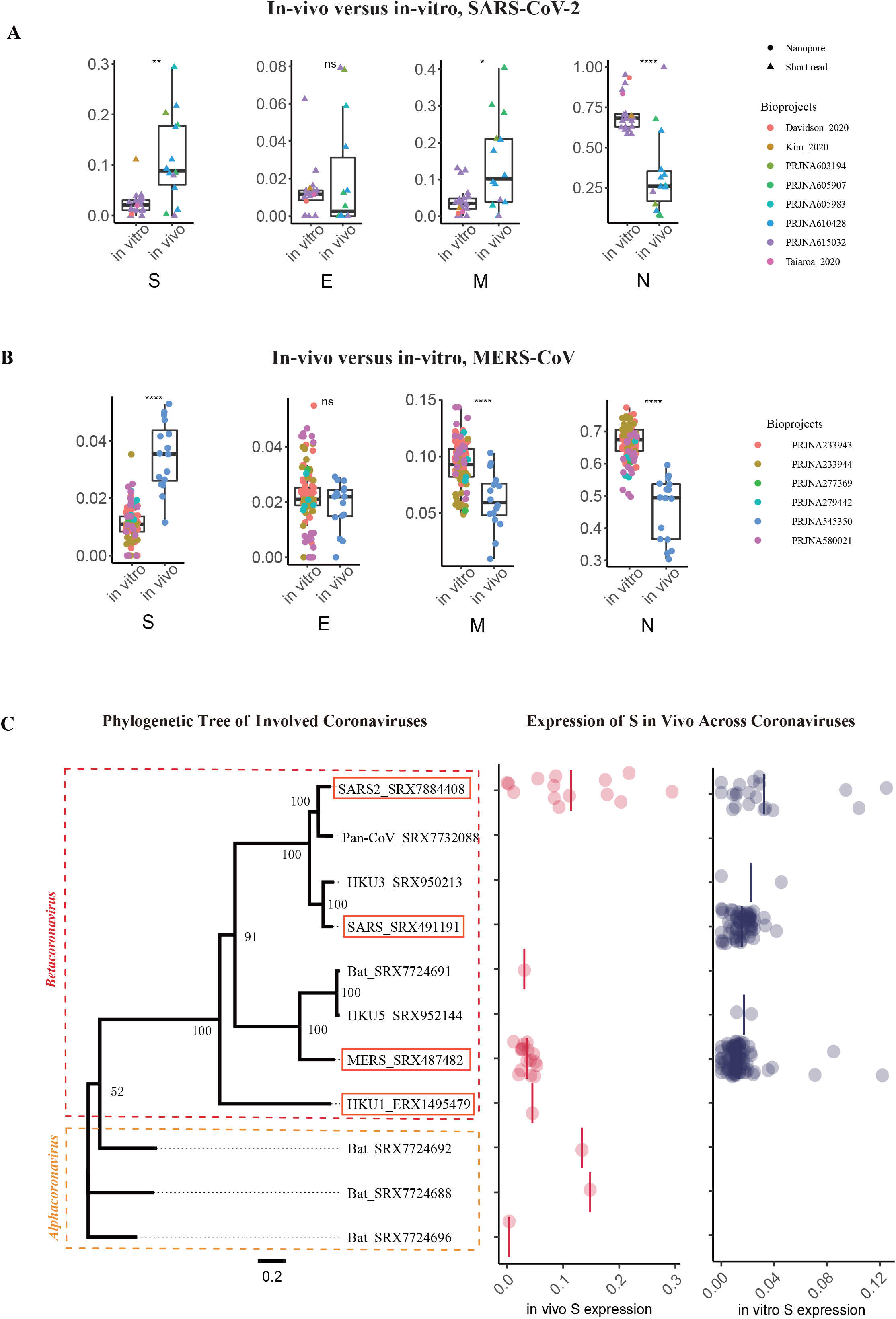
Comparison of *in-vivo* and *in-vitro* sgRNA expression. **(A) and (B)** Expression profile of SARS-CoV-2 and MERS-CoV, both *in-vivo* and *in-vitro* datasets were included. It should be noticed that two third generation sequencing technology data were added as complementary datasets to SARS-CoV-2 *in-vitro* plot, a math model was applied to adjust long read expression ratio into an adapted version which was comparable with short read archive datasets. Interestingly, higher levels of S and M expression ratio and lower level N expression ratio were observed in *in-vivo* sample versus *in-vitro* sample in these two coronaviruses. **(C)** Phylogenetic tree of involved coronaviruses (left), scale bar indicates phylogenetic distance which were calculated as the ratio of nonidentical base positions to all base positions, taxonomic classification at genus level were indicated at left part. Expression ratio of Spike (S) genes in vivo and in vitro in different coronaviruses (right), each dot represent a biosample, black bars indicate average expression level of responsive virus.

To obtain a clearer perspective on how the relative abundance of SARS-CoV-2 sgRNAs compares to other coronaviruses *in-vivo* and *in-vitro* as well as determine if additional novel sgRNAs have been overlooked, CARONTATOR was utilized to analyze additional coronaviruses. This analysis included OC43, NL63, HKU1 as well as bat and pangolin viruses with high sequence homology to SARS-COV-2 [9, 21] (**Fig 4C**, **Table S1 and Fig. S2**). Some datasets did not yield enough breakpoint reads to be informative. For example, analysis of the the bat virus RaTG13, with the highest homology to SARS-COV-2, yielded only 1 break point read and was therefore omitted from **Fig. 4C**.

Of the different coronaviruses profiled, SARS-COV-2 stands out as having the highest levels of S sgRNAs, especially *in-vivo* (see discussion below and **Fig. 4C**). Our analysis indicates that this is independent of viral strain as it is present at high levels in different strains identified *in-vivo* (**Fig. 2**). The high levels of Spike protein may play a role in the viruses ability to cross the species barrier (see discussion below) and it’s high rate of infectivity. In agreement, we noted that the relative levels of the Spike sgRNA is positively correlated with coronavirus infectivity. Viral infectivity and levels of S sgRNAs ***in-vivo*** are as follows: SARS-COV-2> HKU1> MERS [28]. However, S protein levels alone are not sufficient to cause high levels of SARS-CoV-2 transmissibility, as factors such as Spike protein stability, receptor aviditiy [29] and viron stability[30], also contribute to viral transmissibility.

### Mutations in the RTC reverse the expression of N and S sgRNAs in vitro and in-vivo

We also observed mutations in viral RTC components that altered the expression profile of S to N. Specifically, the viral strain Kim 2020 had one unique non-synonymous mutation in the RTC component nsp3, a papin protease that binds the N and M protein (**Fig. 3**). The transcriptome generated *in-vitro* for this viral strain showed a dramatic increase in the S to N ratio, mimicking the expression profile of viruses found *in-vivo* (**Fig. 3**, **Fig. S6**). Interestingly, a viral strain identified *in-vivo* (SRX7852918), had two non-synonymous mutations in nsp3, as well as nsp6 and nsp12 and had an *in-vitro* like transcription profile, with a decreased S to N ratio (**Fig. S6**).

The observation that mutations in nsp3 occur in the two viruses with altered gene expression is thought provoking. Nsp3 is reported to to bind TRS’s, the 3’ end of the viral genome, the global viral RNA packaging signal as well as the N and M proteins [23, 31, 32]. Additionally, phosphorylation of the N protein has been reported to alter it’s conformation to preferentially bind viral RNA and as mentioned above for IBV, promote TRS readthrough during the generation of long sgRNAs [26, 33]. This observation tentatively implies that mutations within nsp3 affect the relative abundance of sgRNAs by acting in a global mechanism that influences overall viral structure and may act in concert with the mechanism described above for IBV. Additionally, the altered relative abundance of N sgRNAs *in-vivo* and *in-vitro* due to mechanisms discussed above, may feedback on it’s interaction with Nsp3 and influence the function of mutations in Nsp3 *in-vivo* and *in-vitro* (**Fig. 4A-B**).

## DISCUSSION

The vast amount of sequence data generated for SARS-CoV-2 thus far has primarily been used for the typing and following of emerging viral strains. Although this is important, we felt such a focus could be an under-utilization of a valuable information. By developing the Coronatator informatics pipeline, we took a step beyond the characterization of viral strains and described coronavirus viral sgRNA expression and uncovered novel and conserved sgRNAs with unknown function that are generated via a non-canonoical TRS pairing mechanism (**Fig. 2**). Functional prediction for some of these novel putative proteins is still ongoing. We tentatively show that a homolog of SARS-CoV-2 pORF2b in pangolin virus shares extensive similarity with human IL17RB’s ligand binding domain (**Fig. 2D**). It is curious that a coronaviruse may generate a peptide that could theoretically disrupt IL17B and IL17E (IL25) signaling as they are generally associated with promoting or inhibiting inflammatory responses in specific contexts. Future proteomic studies and/or ribosome sequencing studies will be required to verify the production of the protein products encoded by the novel sgRNAs idenrified here.

The analysis presented here also implicates that different strains of SARS-CoV-2 express sgRNAs at different levels (**Table S3**, **Fig. 3**), especially for the newly discovered sgRNAs. Our findings underscore that a true understanding of viral pathogenesis in terms of sgRNA expression can only come from thorough sequencing of patient samples in which the virus is under selective pressure. This begs for in-depth case examination, in which thorough sequencing and analysis is conducted for different stages of COVID-19 on a strain by strain basis. This would result in truly individualized patient care.

Although other zoonotic viruses may share extensive sequence similiarity to SARS-CoV-2 at the gene or genomic level, similarity alone is not sufficient for the generation of pathogenic human viruses. Generally not considered during discussion of zoonotic viral orgins, the specific expression level of viral genes, such as the Spike protein, are likely important for crossing the species barrier. For example, considering the vast number of un-sampled zoonotic viruses, it is likely Spike proteins capable of crossing the species barrier already exist, yet are not expressed at sufficiently high levels to enable sustainable inter-human transmission. However, low level Spike protein expression would allow sporadic transmision from bat to human, yet would not be sustainable as human to human transmission would be low due to low S protein expression as well sanitary enivronments that do not exist for bats. In agreement, it has been observed that people living in proximity to bat caves harbor virus specific antibody without ever experienceing severe disease [34].

Our analysis of the meta-transcriptomic data sets identified numerous sources of RNA, such as host RNA as well as microbial RNA (although not optimally captured). In a time when it is unclear why some people succumb to SARS-CoV-2 infection while others do not, these valuable sequences should not be wasted and could be made more useful if more clinical information is shared for these data sets. Most GISAID entries for SARS-CoV-2 have a meta-transcriptomic dataset that supports it. However, current GISAID entries that simply outline the viral genome sequence and strain far out-number the raw read entries we identified in SRA. Sharing the raw read information will greatly help researchers study this virus and ultimately curb it.

## CONCLUSIONS

We developed a bioinformatics pipeline CORONATATOR that can take meta-transcriptomic sequencing reads generated from coronavirus samples and analyze the sub-genomic RNA profiles of the underlying virus, akeen to a transcriptome for the virus. For emergent viruses, as in the case of SARS-CoV2, homology search was usually the first and only choice of predicting viral ORFs after sequencing was done. Now our tool can provide additional evidence. By applying it to large number of SARS-CoV2 and related viral datasets, interesting biology about these coronaviruses were revealed. In addition to define core and predict novel ORFs, our results suggested, for beta-coronaviruses, the spike to nucleocapsid ratio to be a potential tunable in adjusting viral life style and the elevation of this ratio in SARS-CoV2 may contribute to its strong transmissibility. The methods and findings presented here provides a valuable resource for future genomic studies of coronaviruses.

## METHODS

### Data collection

All sequencing data used were collected from NCBI Short Reads Archive (SRA). Some nanopore datasets were downloaded from online repository described in their respective manuscripts [10]. The bioprojects were located by searching with key words “coronavirus” and with manual curation, only meta-transcriptomic data were kept Raw reads files were downloaded from SRA using wget with a customized script, SRAtoolkit were used to generate compressed fastq files from downloaded sra files. After initial sequence alignment using bwa with reference genome sequences of SARS-CoV, SARS-CoV-2 or MERS-CoV, samples with too few viral reads were filtered out. CORONATATOR only uses reads generated from second generation technologies (Illumina), nanopore data were used for comparison.

### Coronatator

CORONATATOR were a series of perl and bash scripts developed for profiling and analysis of RNA-Seq data from coronavirus. It consists of 3 major steps, including preprocessing, breakpoint identification, sgRNA calling and profiling, details below.

### Preprocessing

BAM files were generated from sequence alignment with reference genomes of SARS-CoV, SARS-CoV-2 or MERS-CoV, for viruses from bat and pangolin, responsive genome assemblies were obtained from NCBI as references. SNPs were called and filtered with bcftools [35] and annotated with vcf-annotator [36]. In addition, consensus genome sequences were also generated with filtered SNPs for further analysis.

### Breakpoint identification

Breakpoints were identified from alignments with soft or hard clips, these alignments were all partial alignments largely caused by reads with recombination joints, which was generated by the mechanism through which coronavirus produce their sgRNA. In this step, a matrix of reads’ information, breakpoint sites, CIGAR strings together with possible TRS sequences was generated.

### sgRNA calling and profiling

Typical sgRNAs were identified and defined by two breakpoint coordinates on a reference genome sequence, these sites were obtained by extracting breakpoints from partial alignments, i.e. one from primary alignment and the other from supplementary alignment. To recognize possible TRS pattern, sequences between breakpoint pairs were extracted from previous generated consensus genome sequences. After that, corresponding genes of called sgRNAs were identified by manually comparing the distances between start codons of known viral genes and their breakpoints. Biosamples with more than 20 sgRNAs were used for further analysis, in these samples, sgRNAs were counted by genes and normalized by total sgRNA count to obtain a transcription profile matrix.

### Novel ORF identification

Potential ORFs were predicted using Prodigal [37] with -s arguments to write all potential genes. An in-house python script was also used to identify very short ORFs. Then for sgRNAs with multiple bioproject support, we calculated and sorted the distances between their breakpoints and all identified start codon sites. ORFs that start closest to upstream breakpoints were bookmarked and manually checked for verification.

### Sequence alignment and phylogenetic analysis

Consensus genome sequences of SARS-CoV-2, SARS-CoV, MERS-CoV and biosamples from bat or pangolin or other human coronavirus with more than 20 sgRNAs were used for phylogenetic analysis. Multi-sequence alignment were performed with MAFFT [38], Maximum likelihood consensus trees were constructed using IQ-TREE [39] with 1000 bootstrap times.

### Converting Nanopore sgRNA proportion to short reads’

Kim et al included both nanopore data and short read data. The ratios between the two were used to convert the other nanopore data sets to proportions comparable with others in this study.

### Plots and statistical analysis

Heatmaps showing gene expression profile were produced using ‘heatmap.plus’ package. SgRNA expression dot plots and boxplots were made with ‘ggplot2’ package to compare difference between gene expression among different sample origin, T-test and wilcoxon test were used for statistical analysis.

### Function annotation

Novel peptide sequences were aligned with EMBL online tool FASTA (https://www.ebi.ac.uk/Tools/sss/fasta/) against UniProtKB/Swiss-Prot database with default arguments. NCBI CD Blast online service was used to identify protein domains.

### Sequence conservation

To check for sequence conservation of putative peptides in related viral species, we generated a reference database containing all predicted ORFs from related viral genomes. DC MegaBlast (DisContinuous MegaBlast) was used to search for inter-species homologs. Arguments were set as follows: window_size 0, gapopen 0, gapextend 2, penalty −1, reward 1, num_alignments 1. A group of homologous ORFs were then subjected to multiple sequence alignment (MSA) using MAFFT. After that CLUSTAO (Clustal Omega) was used to calculate an identity matrix for the MSA result. The same procedure was performed for both nucleotide and amino acid sequences.

## ABBREVIATIONS

sgRNA: sub-genomic RNA
ORF: open reading frame
TRS: Transcription Regulatory Sequences
COVID: Corona virus disease
MERS: Middle Eastern Respiratory Syndrome
SARS: Severe Acute Respiratory Syndrome
CORONATATOR: CORONAvirus annoTATOR

## DECLARATIONS

### Ethics approval and consent to participate

Not applicable.

### Consent for publication

Not applicable.

### Availability of Data and Materials

All used sequenceing data are accessible with accession number provided in supplementary table 1, code of CORONATATOR is accessible at: https://github.com/15274972986/CORONATATOR.

### Authors’ contributions

Conceptualization: Lei Chen; Methodology: Lei Chen, Lyu Lin; Investigation: Lyu Lin, Ru Feng, Mingnan Zhang, Yinjing Liao; Visualization: Lyu Lin, Qiyu Gong; Supervision: Lei Chen, Xiaokui Guo, Bing Su, Yanjiao Zhou; Writing—original draft: Lei Chen, Yair Dorsett, Lyu Lin; Writing—review & editing: Lei Chen, Yair Dorsett, Lyu Lin, Ru Feng.

## Acknowledgements

We thank Dr. Qiming Liang of Shanghai Institute of Immunology for his insightful suggestion.

## SUPPLEMENTARY INFORMATION

**Additional file 1: Fig. S1:**
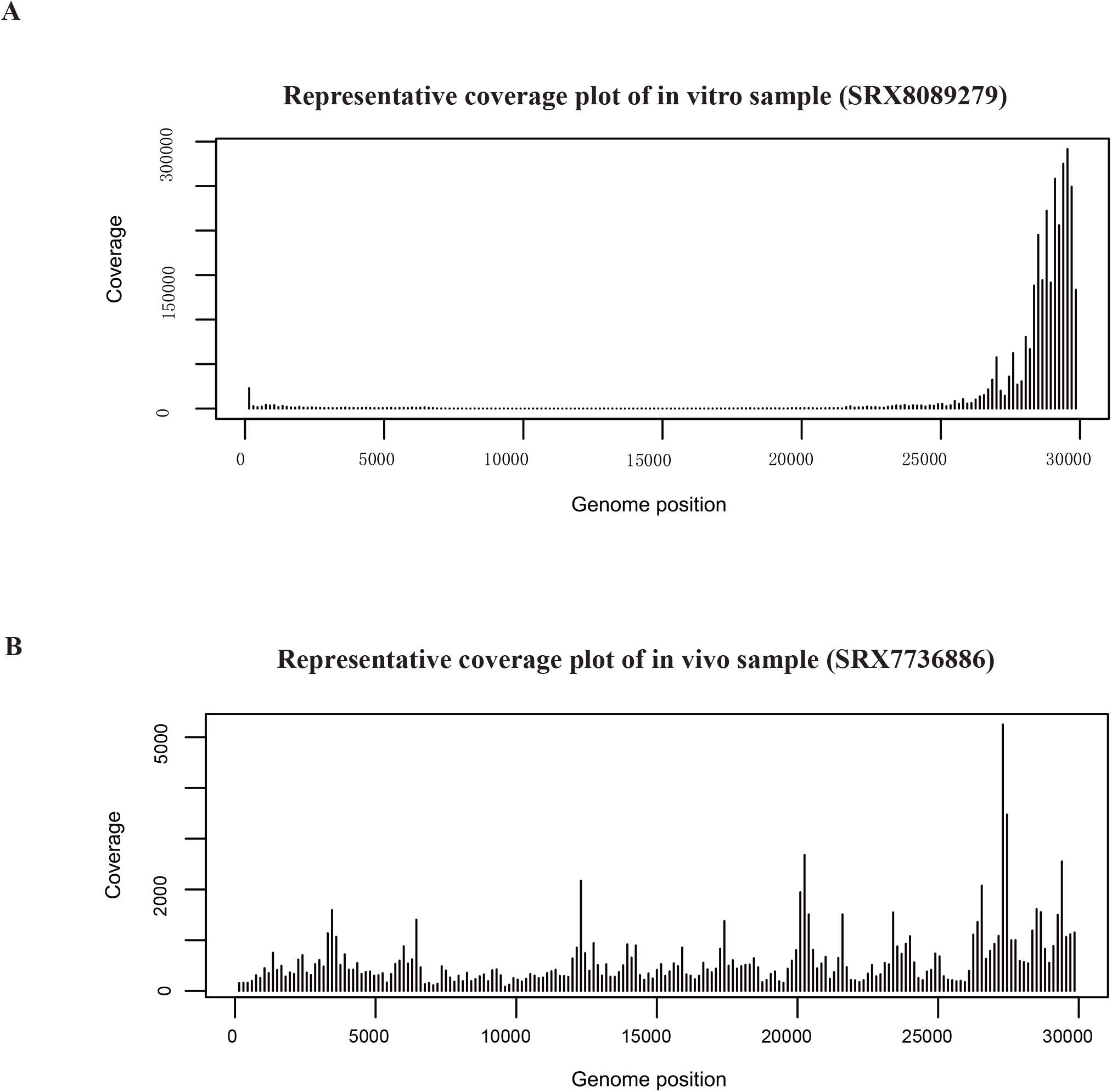
Coverage plot for in-vivo and in-vitro datasets. **(A)** Coverage plot for SRX8089279, which is a representative of in-vitro sample. In-vivo RNA-Seq reads relatively evenly mapped to viral genome, indicating a genomic RNA dominated sample. **(B)** Coverage plot for SRX7736886, which is a representative of in-vivo sample. In-vitro reads resulted a dense mapping at 3’ and 5’ end of the genome, which revealed active viral transcription and replication in cultured cells.

**Additional file 2: Fig. S2 :**
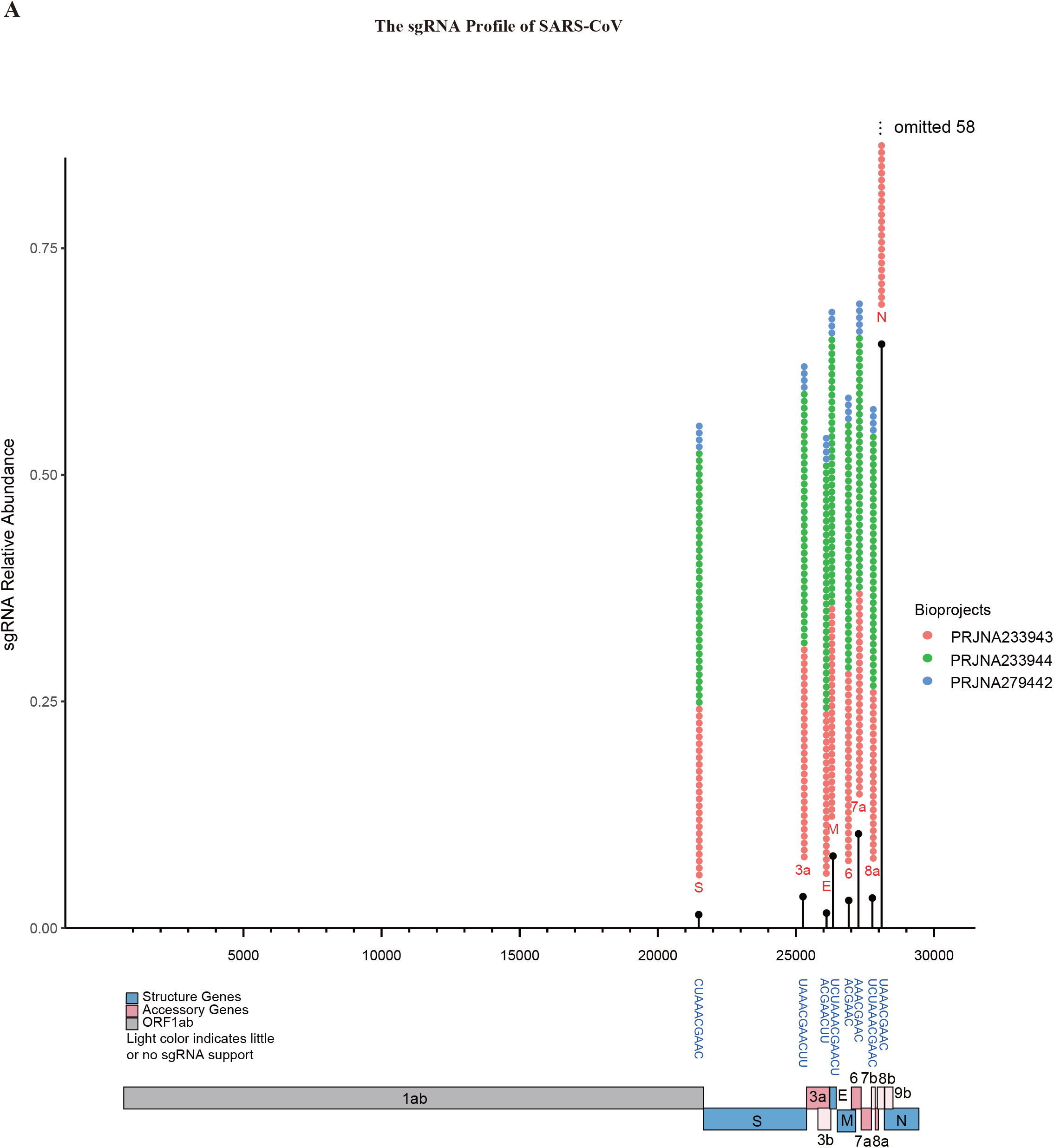
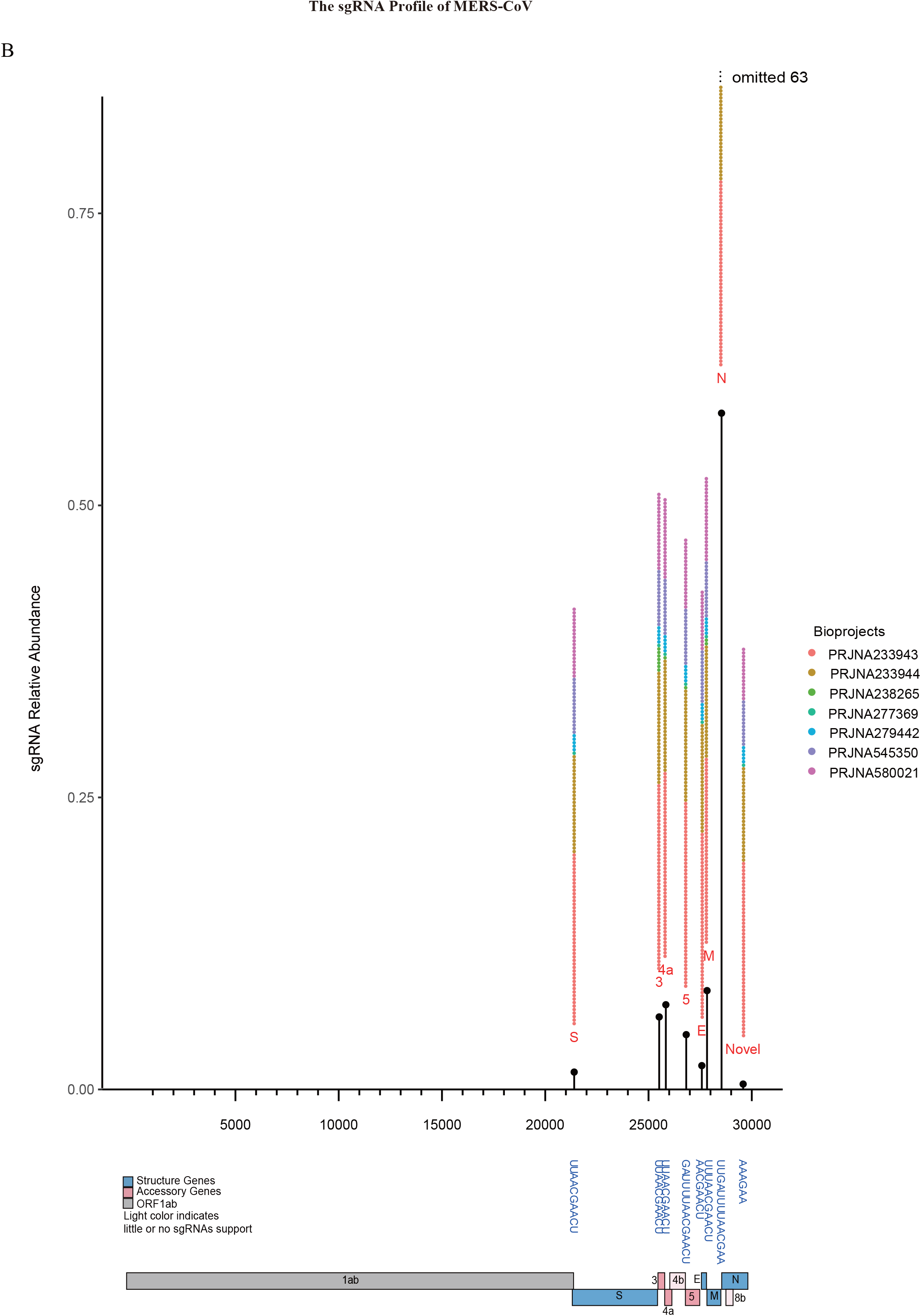
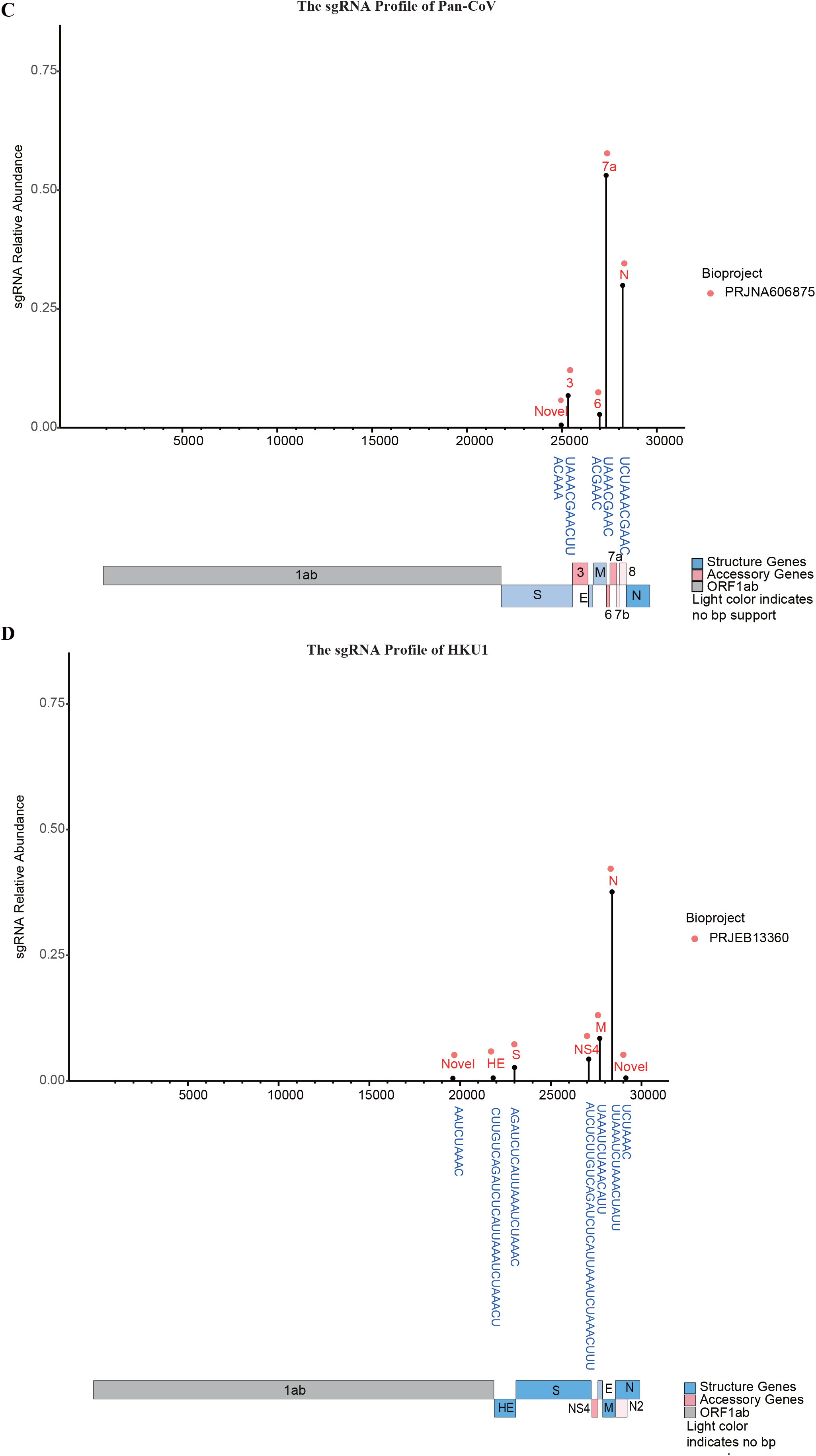
Breakpoint profile of 4 coronaviruses. **(A)** Breakpoints profile of SARS-CoV; **(B)** Breakpoints profile of MERS-CoV; **(C)** Breakpoints profile of Pangolin-CoV; **(D)** Breakpoints profile of HKU1, interestingly, this virus uses long TRS for discontinuous sgRNA production.

**Additional file 3: Fig. S3:**
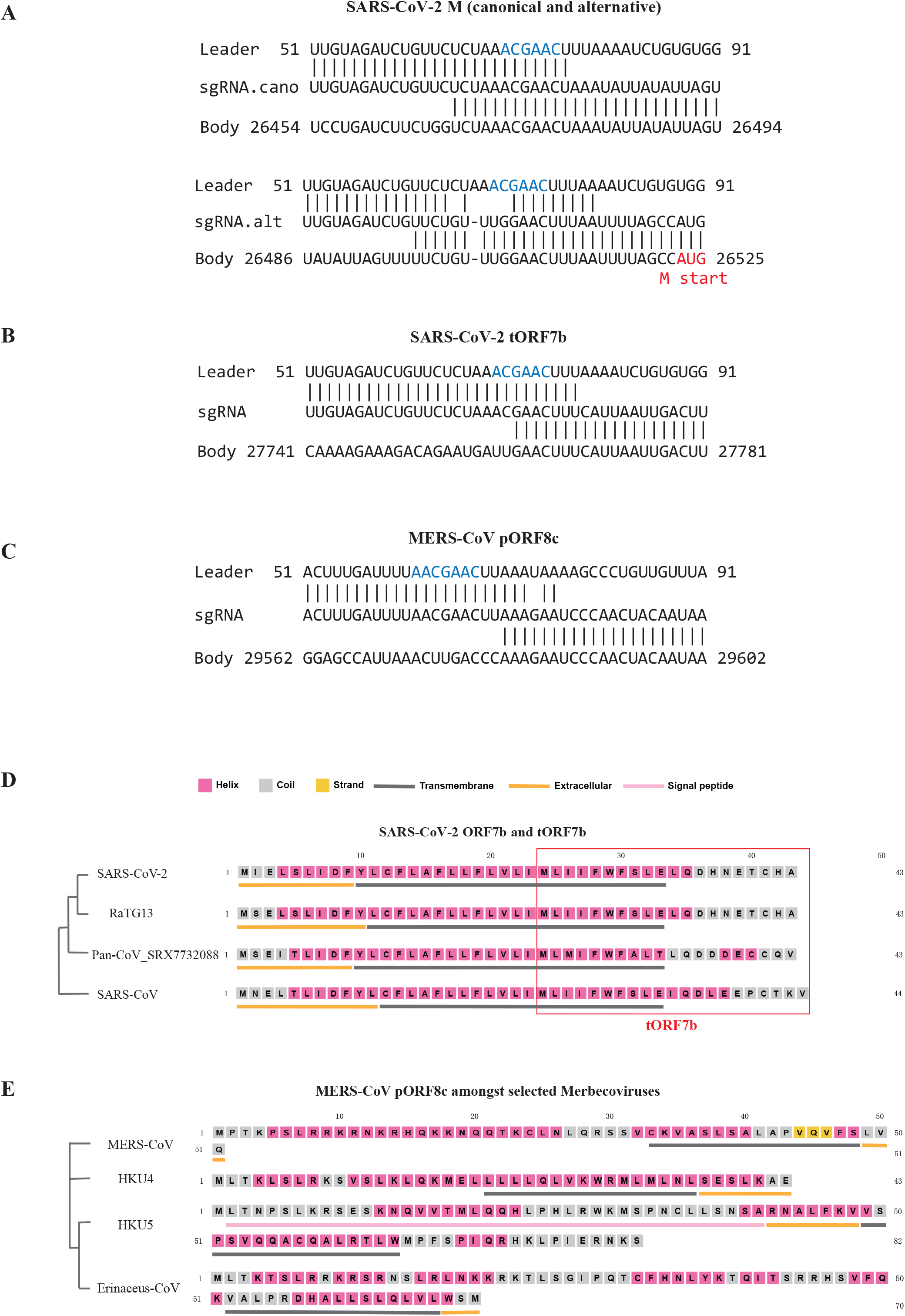
Novel sgRNA breakpoint and TRS sequence. **(A)** Alternative TRS of M in SARS-CoV-2. The canonical TRS region (upper panel) has 12 bases while the novel one has only 6 (lower panel), start codon of M was shown in red. **(B)** TRS for tORF7b in SARS-CoV-2, which has 7 bases overlapped with leader TRS. **(C)** TRS for pORF8c in MERS-CoV. **(D)** and **(E)** Structural conservation of novel peptide translated from newly discovered sgRNA truncated ORF7b and putative ORF8c, left panel demonstrates consensus phylogenetic tree of responsive coronavirus determined by genomic sequences, right panel compared second structure of novel peptide predicted by PSIPRED Workbench.

**Additional file 4: Fig. S4:**
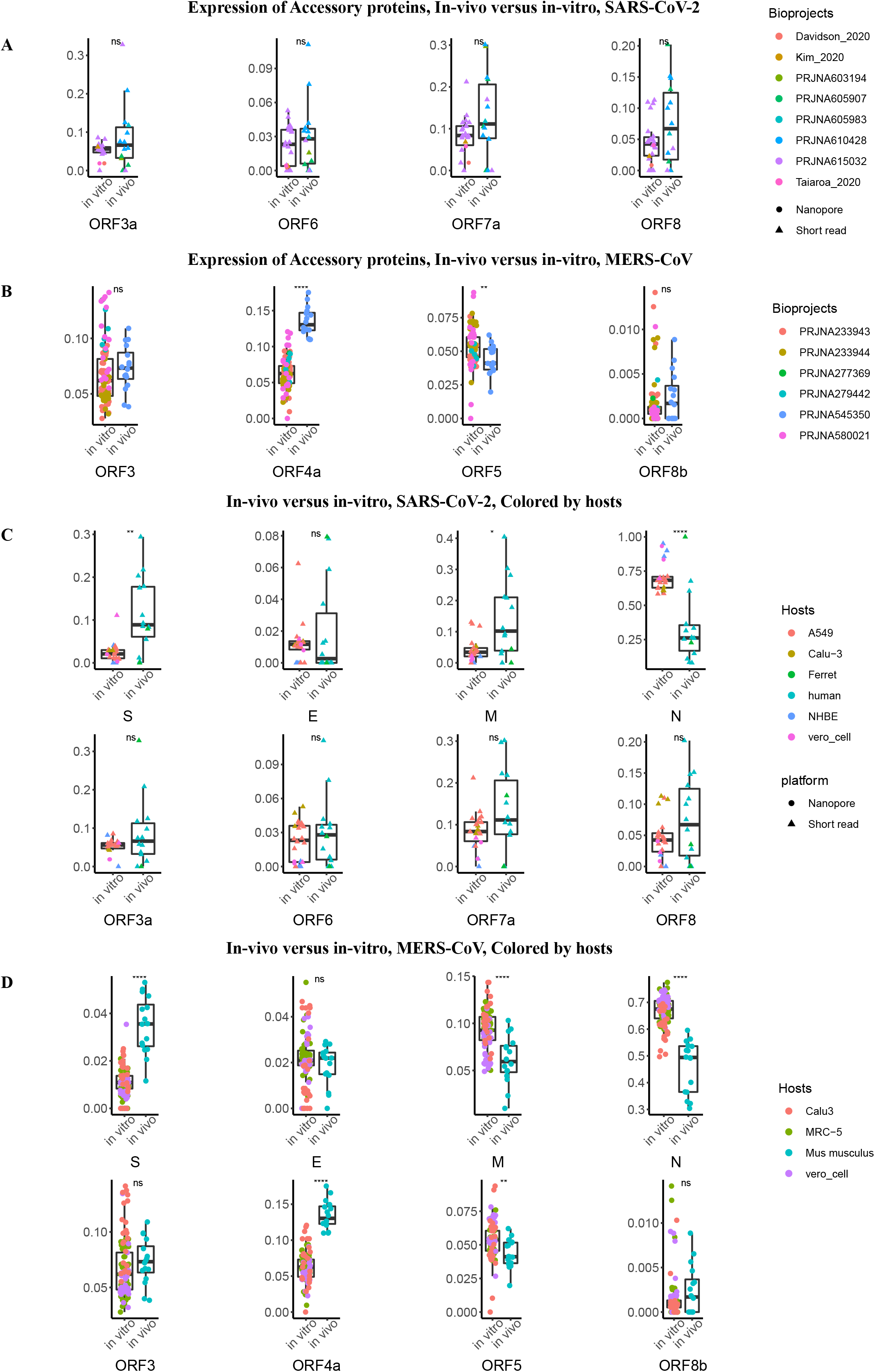
Detailed gene expression profile of SARS-CoV-2 and MERS-CoV. **(A)** and **(B)** Detailed accessory gene expression profile of SARS-CoV-2 and MERS-CoV, between in-vivo and iv-vitro datasets. Remarkably, MERS-CoV had higher ORF4a in-vivo expression level while lower in-vivo ORF5 expression level. **(C)** and **(D)** Gene expression level of SARS-CoV-2 and MERS-CoV among different hosts.

**Additional file 5: Fig. S5:**
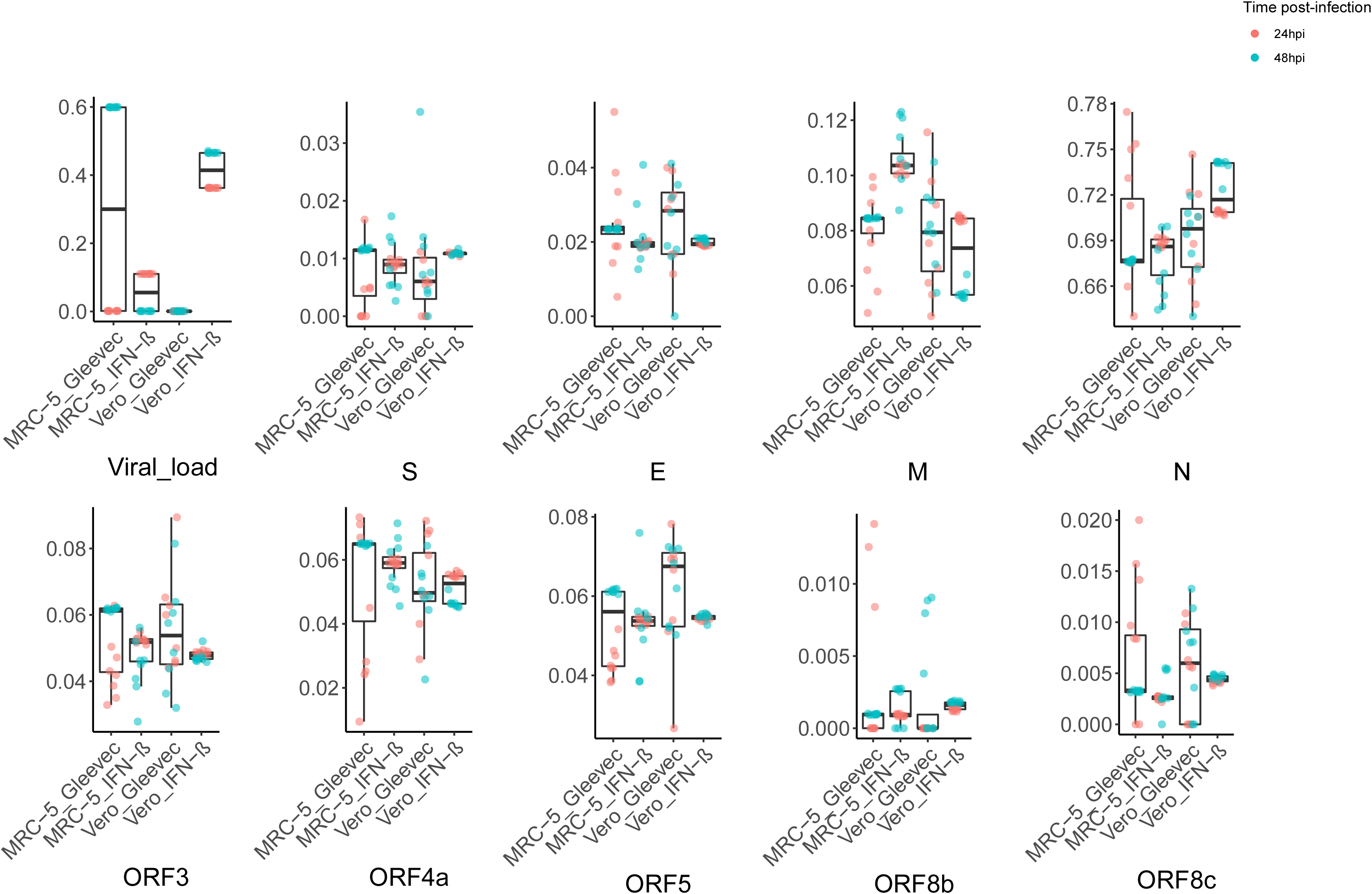
Detailed gene expression profile of MERS-CoV in PRJNA233943 & PRJNA233944.In this study, cells infected with MERS-CoV were treated with different drugs, i.e. Gleevec and IFN-β, after 24 or 48h post-infection, distinct pattern can be observed from viral gene expression profile as condition alters, especially for IFN-β, treated 24 hpi and 48 hpi resulted in distinct expression levels among several structural and accessory genes. It also indicates that our analytical pipeline CORONATATOR is a powerful and sensitive tool for analyzing how experimental manipulation effects the relative expression of specific sgRNAs.

**Additional file 6: Fig. S6 :**
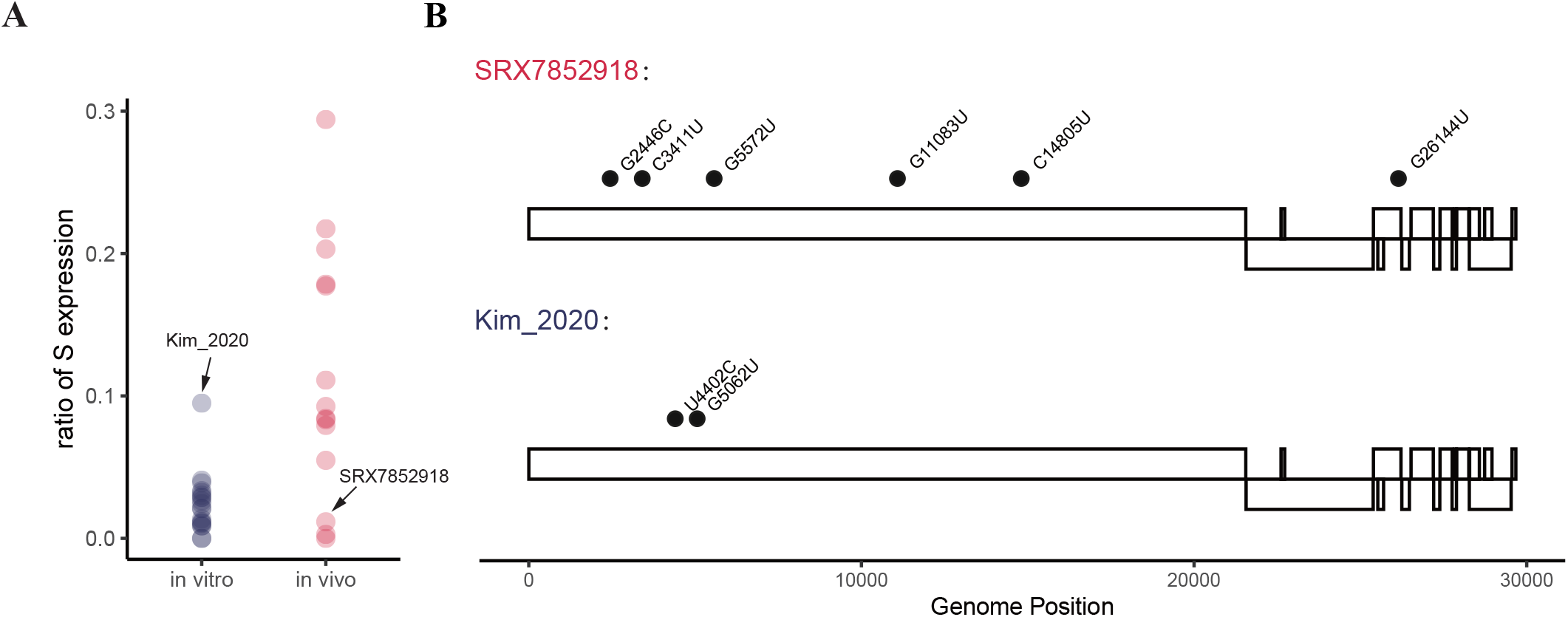
Outliers in expression profiles habour interesting SNPs. **(A)** and **(B)** The in-vivo sample from Kim et al 2020 had S expression level similar to that of in-vitro samples. While also have a few mutations in ORF1a that’s not found in other viral strains. Mirroring this, the in-vitro sample SRX7852918 have S expression level similar to that of in-vivo samples, and hold several private mutations in ORF1a as well.

**Additional file 7: Table S1 :** Meta information of samples collected.

**Additional file 8: Table S2 :** Annotation of SARS-CoV-2, SARS-CoV and MERS-CoV.

**Additional file 9: Table S3:** SARS-CoV-2 sgRNA abundance across samples.

**Additional file 10: Table S4:** Conservation of selected novel proteins.

**Additional file 11: Table S5:** List of novel sgRNAs.

